# Insights into early stages of the establishment of host:algal endosymbioses: Genetic responses to live versus heat-killed algae and bacterial prey in a sponge host

**DOI:** 10.1101/2021.06.16.448416

**Authors:** Sara Camilli, Vasiliki Koutsouveli, Chelsea Hall, Lillian Chang, Oriol Sacristan-Soriano, Malcolm Hill, Ana Riesgo, April Hill

## Abstract

The freshwater sponge *Ephydatia muelleri* and its *Chlorella-*like green algal partner is an emerging model for studying animal:algal endosymbiosis. The sponge host is a highly tractable laboratory organism, and the symbiotic algae are easily cultured. We took advantage of these traits to experimentally interrogate fundamental questions about cellular mechanisms that govern the establishment of durable intracellular partnerships between hosts and symbionts in facultative symbioses. We modified a classical experimental approach to discern the phagocytotic mechanisms that might be co-opted to permit persistent infections, and identified genes differentially expressed in *E. muelleri* sponges early in the establishment of endosymbiosis. We exposed algal-free *E. muelleri* to live native algal symbionts, native heat-killed algae, and bacteria, and then performed RNASeq so we could compare patterns of gene expression in each treatment. We contrasted differential gene expression patterns between potential food items (bacteria and heat-killed algae) and the live native *Chlorella*-like symbiont. We found a relatively small but interesting suite of genes that are uniquely and differentially expressed in the host exposed to live algal symbionts, and a larger number of genes uniquely triggered by host exposure to heat-killed algae. One of the host genes, an ABC transporter that is downregulated in response to live algal symbionts, was further evaluated for its possible role in establishment of the algal symbiosis. We discuss the gene expression profiles associated with host responses to living algal cells in the context of conditions necessary for long-term residency within host cells by phototrophic symbionts as well as the genetic responses to sponge phagocytosis and immune driven pathways.

## INTRODUCTION

For intracellular symbionts that provide benefits to the host (i.e., mutualists), the host cell represents a potentially dangerous habitat. Hosts can digest or mount immunological defenses against foreign entities (Schwarz 2008; Schmid-Hempel 2009; Davy et al. 2012; Hill and Hill 2012; Hill 2014), even those that provide them with physiological benefits. Despite the challenges of living inside a host cell, intracellular habitats have been repeatedly invaded by a diverse assortment of microbes with varying degrees of specificity and duration of occupancy (e.g., Nowack and Melkonian 2010; Romano et al. 2013). However, the mechanisms that permit stable intracellular occupancy are often poorly understood. Phagocytosis and organellar or cytoplasmic residency expose the symbiont to mortal threats, and each stage of phagocytotic processing presents opportunities for the symbiont to circumvent host cellular processes.

Several pathways of host-cell occupancy have been proposed (e.g., Schwarz 2008; Hill and Hill 2012), but the reality is that we have limited understanding of the precise mechanisms that permit intracellular residency. We have a great deal to learn about the cellular and genetic mechanisms that symbionts subvert to avoid digestion or destruction. The work described here represents an effort to gain important perspectives at a fine scale genetic resolution on these ecologically essential symbiotic interactions. We compare host responses to food/prey items, heat-killed symbionts, and live symbionts to discern genetic pathways that may be essential for stable symbioses. Mutualistic intracellular phototroph:heterotroph symbioses are essential features of many ecosystems. The most well studied animal:algal symbionts involve members of the Symbiodiniaceae in marine environments and *Chlorella*-like green algae in freshwater systems (Venn, Loram & Douglass 2008). The Symbiodiniaceae include dinoflagellates that form symbioses with an astonishing diversity of mostly invertebrate hosts in tropical and sub-tropical marine habitats (LaJeunesse et al. 2018). Photosynthetically-derived nutrients translocated from the dinoflagellate partner to the host can meet large percentages of the host’s energy budget (e.g., Muscatine 1990; Whitehead & Douglass 2003; Weisz et al. 2010; Davy et al., 2012; Hill 2014; Schlichter 2015), which contributes to the impressive coral growth in shallow oligotrophic tropical waters (Venn, Loram & Douglass 2008). The dinoflagellates appear to receive nutrients from the host, though the intricacies of that exchange are still being elucidated (Davy et al. 2012). A great deal of interest on the initiation of the host:symbiont partnership in this area has focused corals, and the process of horizontal transmission (Ali et al. 2019). The emphasis here is most often at a coarse level: hosts “acquire” symbionts without much focus on the intricate cellular processes that permit invasion of the host and cross-generational stability. However, several studies have provided a foundation for understanding the host genetic response and cellular mechanisms governing these mutualisms (e.g., reviewed in Weis 2019). As with the Symbiodiniaceae symbioses, *Chlorella*-based symbioses are ecologically important features of freshwater ecosystems, and our understanding of aspects of intracellular residency has been improved through detailed study of these symbioses. For instance, the cellular and genetic pathways involved in *Chlorella* infection by *Paramecium bursaria* have begun to be elucidated (e.g., Kodama et al. 2014). *Chlorella* occupy perialgal vacuoles derived from digestive vacuoles in *P. bursaria*, and growth of the host is impacted by algal presence (Karakasian 1963). While *Chlorella* appear to avoid digestion through patterns of spatial distribution within *Paramecium*, and can persist despite lysosomal fusion with the perialgal membrane, important molecular aspects of the infection process remain unclear (Kodama and Fujishima 2005, 2007, 2012; Kodama et al. 2007). Another *Chlorella*-based symbiotic partnership involves freshwater cnidarians who receive photosynthetically-fixed carbon from the algae (Muscatine & Hand, 1958). *Hydra* spp. typically maintain *Chlorella* in their gastrodermal cells in organelles surrounded by a perialgal membrane of host origin. During intracellular residency by the symbiont, *Hydra* hosts engage distinct molecular and cellular pathways, including the upregulation of phosphate transporters and glutamine synthetases (e.g., Ishikawa et al. 2016; Hamada et al. 2018).

In addition to the important symbiotic systems mentioned above, members of the phylum Porifera harbor a diverse assortment of photosynthetic partners (Taylor et al., 2007), including intracellular relationships with members of the Symbiodiniaceae (Hill 1996; Hill et al. 2011; Strehlow et al. 2016; Ramsby et al. 2017) and *Chlorella*-like algae (Hall et al. 2021). We recently introduced the freshwater sponge, *Ephydatia muelleri* and its *Chlorella*-like algal symbiont as a tractable model system to study endosymbiosis (Hall et al. 2021). In this facultative association, both interacting species are easily grown alone and in symbiosis, making it possible to study the mechanisms that lead to successful partnerships in a relationship that is mutually beneficial but not fully interdependent. The exact mechanical process through which symbionts enter and disperse throughout these sponges is still relatively unknown, but they can be taken up by, and divide in, a variety of cells including choanocytes (Wilkinson 1980). Symbionts are found in intracellular compartments as early as 4 hours post algal infection and persist in regions surrounding choanocyte chambers (Hall et al. 2021).

A major goal of the work presented here was to discern pathways that differentiate successful intracellular residency of a potential mutualistic symbiont from digestion of prey/destruction of foreign entities. That is, how does a successful symbiont avoid triggering the host’s digestive response? To answer this question, we employed a classic experimental approach (i.e., we exposed hosts to live and heat killed algae) combined with advanced molecular tools. We sequenced the transcriptomes of sponges recently hatched from gemmules that were exposed to three “foreign” entities: living bacteria (a potential food item), living sponge-derived algae (a potential symbiont), and sponge-derived algal cells that had been heat-killed (a potential food item that might have intact symbiosis-related antigens). Each of these treatments was compared to algal-free sponges that had not been exposed to any external material (“control” aposymbiotic sponges). Patterns of gene expression were compared among treatments. Experiments examining the function of one transcript (ABCC4) that is downregulated in algal infected sponges relative to aposymbiotic sponges are presented. We discuss the implications of this work in light of growing interest in the evolution of specificity between hosts and symbionts and the fundamental and realized niche of phototrophic algae.

## RESULTS & DISCUSSION

Among the multitude of possible types of symbiotic associations, none are more tightly integrated than those that involve intracellularity. This is partly due to the operation of strong and reciprocal selective processes. Our overarching objective is to better understand forces shaping intracellular symbioses involving phototrophic symbionts and heterotrophic hosts. We previously demonstrated that *Chlorella*-like green algal symbionts could be isolated from field collected *Ephydatia muelleri*, successfully cultured, and used to infect algal free “aposymbiotic” *E. muelleri* grown from gemmules (Hall et al, 2021). We also reported on RNASeq experiments comparing aposymbiotic and infected sponges one day after the initiation of symbiosis and showed that a variety of conserved pathways were activated in response to the symbionts (Hall et al, 2021). Here, we both examine patterns of gene expression specific to an earlier stage of infection of native algal symbionts with their freshwater sponge hosts and differentiate between genes expressed in response to potential symbionts versus potential food. To this end, we infected aposymbiotic *E. muelleri* with heat-killed microalgae, bacterial food items, and live microalgae (fig. 1) in order to differentiate the host genetic response to establishment of native symbionts from cellular processes activated when food items are processed or when the host is presented with non-viable algae that may have epitopes of living algal cells. Confocal and electron microscopy confirm that algae and bacteria can be found inside host cells between 1-4 hours after infection (fig. 2) and previous work demonstrates that live algae persist and divide within the host while bacteria and other potential prey are digested (Hall et al, 2021).

**Figure 1:**
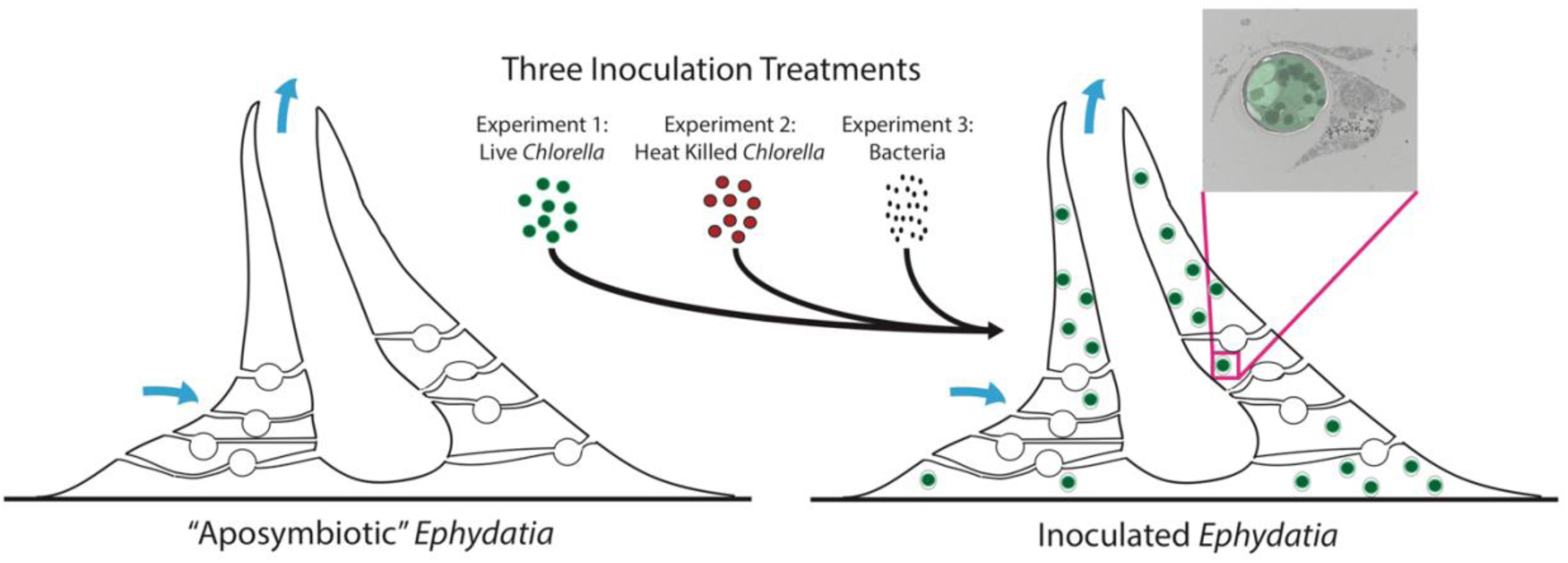
Schematic representation of infection and feeding experiments. Biological triplicate treatments of algal-free “aposymbiotic” sponged inoculated with live algae, heat-killed algae, or bacteria. After Hall et al., 2021.

**Figure 2:**
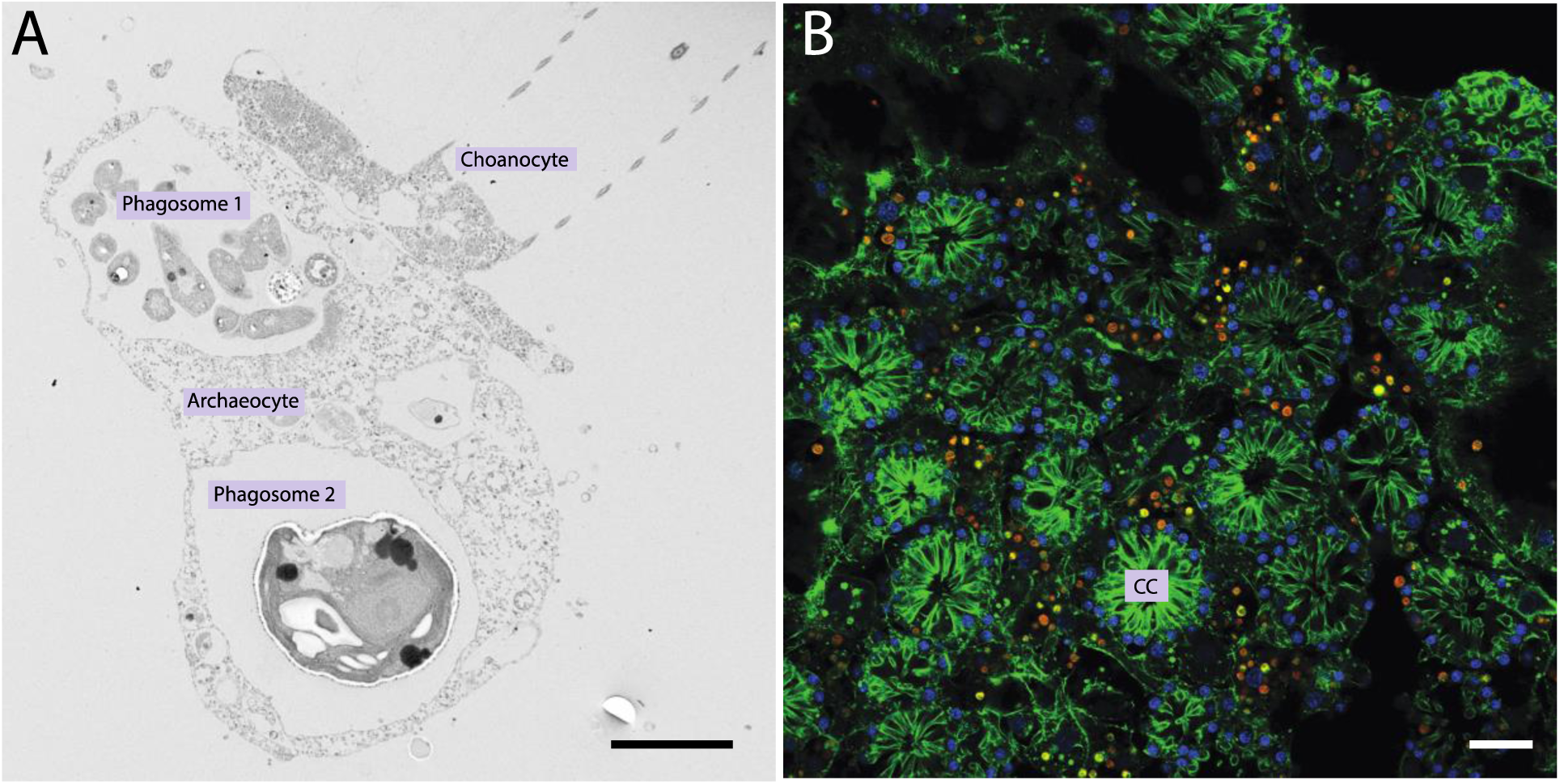
Electron and Confocal Microscopy of *E. muelleri* cells 4 hours post inoculation. A. Transmission electron microscopy image of sponge choanocyte cell abutting an archeocyte cell containing two phagosomes. Phagosome 1 contains multiple bacteria and Phagosome 2 contains one algal cell. Scale bar 2 µm. B. Confocal image of sponge choanocyte region showing intracellular algal symbionts (red) in and around choanocytes (CC, choanocyte chamber). Image shows DNA in blue, F-actin in green, and autofluorescence of algal cells in red. Scale bar 30 µm.

### RNA Sequencing and Assembly

We obtained a total of 331.4 million raw reads, and half were retained during the filtering process (over 143 million reads, see supplementary table S1). For two samples (one in the bacterial fed - BAC and one in the living algal fed - CHLO treatments), the sequencing was not optimal and they were both discarded for the subsequent gene expression analysis. Between 50 and 60% of the reads for each sample were mapped unequivocally to the reference transcriptome (supplementary table S1). The assembly contained 190.96 Mb and 218,986 transcripts and 117,619 genes, with an N50 of 1,527, and a GC content of 45.99% (supplementary table S1). The completeness of the transcriptome was quite high, with over 97% of eukaryotic and over 90% of metazoan conserved genes recovered (supplementary table S1).

Around 32% of the reference transcriptome obtained a blast hit against *refseq* (supplementary table S1), and as expected, the reference transcriptome had bacterial, algal, and sponge genes represented (fig. 3). The majority of genes were of sponge origin (88.9%), though we also observed transcripts representing bacterial genes from the *E. coli* used in feeding treatments, the bacterial symbionts carried by the sponge gemmules used in the experiment, and algal genes from the native algal symbiont. The bacterial and algal genes represented 5.4% of the reference transcriptome. A very small portion of the genes were assigned to other organisms, like viruses or protists (5.7%). Almost all sequences with a blast hit (87%) could be further annotated with Blast2GOPRO (supplementary table S1).

**Figure 3:**
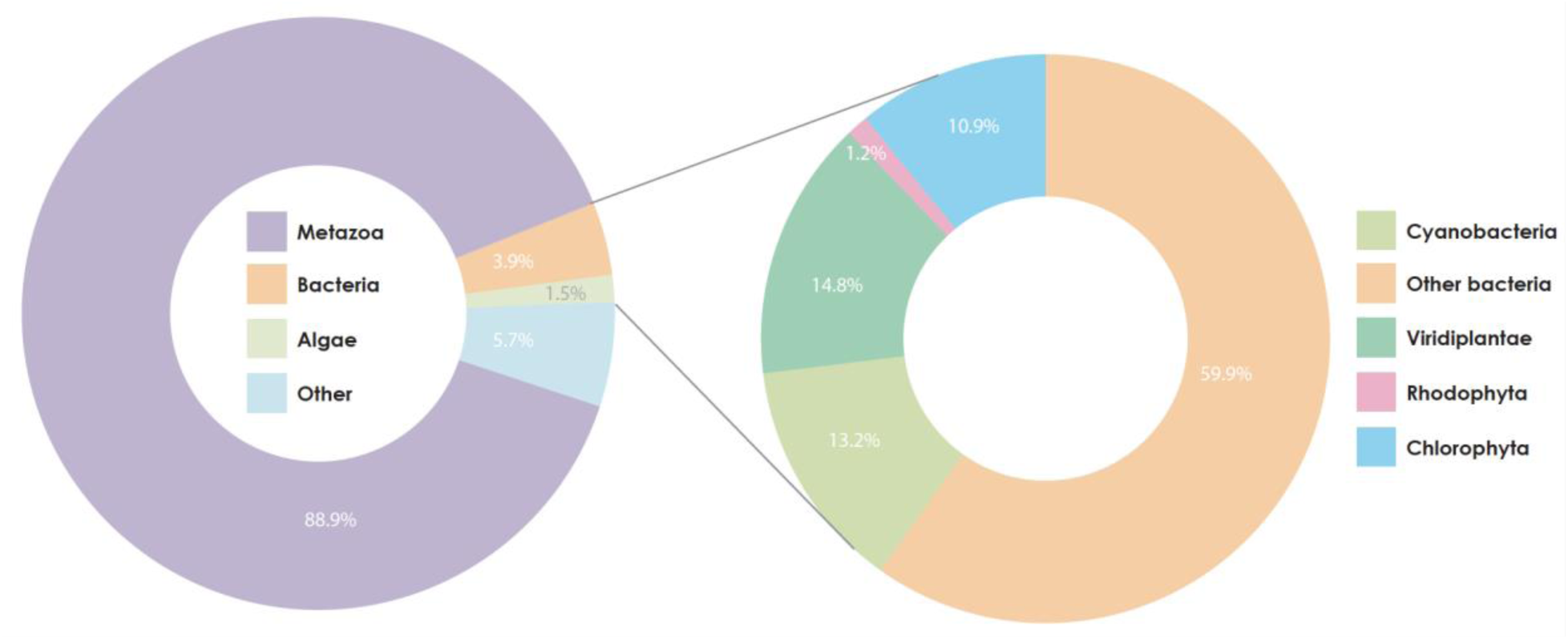
Relative proportions of transcripts in reference transcriptome. The majority of genes represented in the transcriptome are animal in origin. Bacterial genes come from both the *E. coli* that was used in the bacterial infection as well as from bacterial symbionts carried by the sponge gemmules. The Viridiplantae, Rhodophyta, and Chlorophyta genes are from the algal symbiont.

### Putative Symbiosis Related Genes Involved in Stable Residency

The nature of the association between freshwater sponges and *Chlorella-*like algae is facultative, as is the case in many phototroph:heterotroph partnerships, in that both the host and symbiont have the capacity (and in fact do) live some portion of their life cycles separate from one another. When they do associate, it is unclear how the phagotrophic sponge differentiates potential mutualists from potential prey. Here, differential gene expression analysis revealed changes in gene expression between control algal-free (“aposymbiotic”) sponges and each experimental treatment (i.e., bacteria-fed (BAC), heat-killed algae fed (HK–CHLO), live native algae infection (CHLO)) at 4 hours post inoculation (fig. 4), with most genes differentially regulated between the APOc or BAC treatments when compared to either of the algal feeding treatments (supplementary figures S1–S6). Indeed, given the fact that sponge gemmules carry associated bacteria that can be digested as food, the gene expression profile for sponges fed bacteria versus the aposymbiotic sponges contributed to the observation that these sponges had very similar expression patterns (supplementary figure S7) and therefore very few genes were significantly differentially expressed between these two treatments (i.e., only 24 genes showed expression differences between aposymbiotic and bacteria fed sponges, supplementary table S2). Thereafter, given the stable expression profiles between bacteria fed sponges and aposymbiotic sponges (fig. 4 and supplementary figure S7), we subsequently treated these conditions similarly when comparing gene expression with sponges fed live and dead algae.

**Figure 4:**
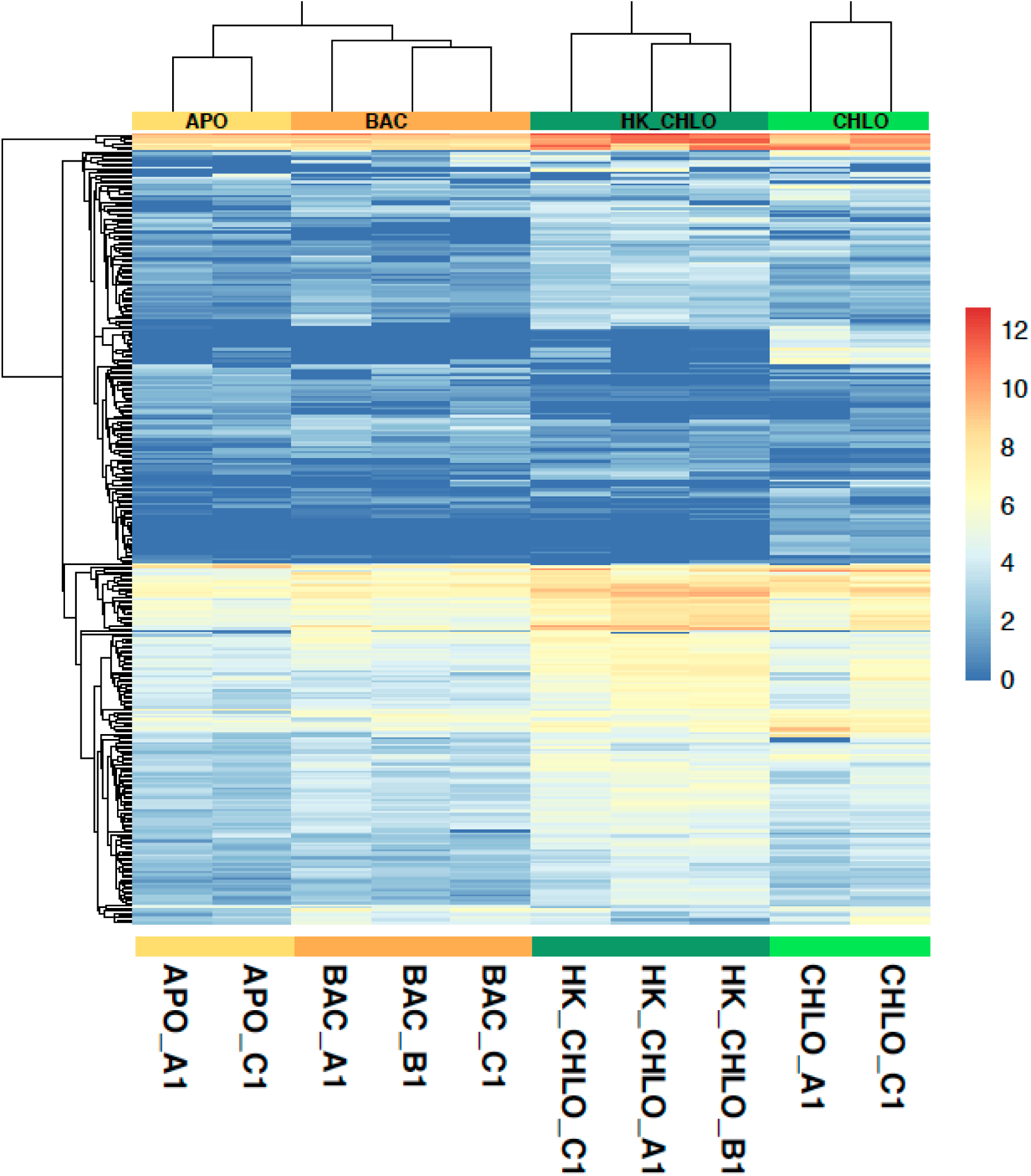
Heat map of differential expression. Heatmap shows all significantly (minimum fold change of 2 and a *p*-value of 0.001) differentially expressed genes across four treatments. Each row of the heatmap represents the z-score transformed log 2 (1 + FPKM) values of one differentially expressed gene across all samples (blue, low expression; red expression, high expression). Aposymbiotic - APO; Bacterial fed - BAC; Heat-killed algal fed - HK-CHLO, and live algal infected - CHLO.

Only around 30% of the genes that were uniquely upregulated in sponges infected with live native symbionts (CHLO) or in those fed heat-killed symbionts (HK–CHLO) were characterized proteins with a known function (fig. 5), the rest being either uncharacterized proteins or putative sponge-specific proteins of unknown function (supplementary table S2). It should be noted that 74% of *E. muelleri* proteins are known to have similarity to other organisms and half of all *E. muelleri* genes can be assigned full functional annotations using BLAST2GO (Kenny et al., 2020). The symbiosis specific genes here are thus enriched regarding the smaller portion of the freshwater sponge genome that is uncharacterized at the protein level. Thus, our data can only describe a subset of the genes that likely play roles in the regulation of stable symbioses until further work on this large body of uncharacterized protein-encoding genes moves forward. Most of the genes upregulated in the in sponges infected with live native symbionts (CHLO) and in those fed heat-killed symbionts (HK–CHLO) were involved in vesicle-mediated transport, endocytosis, protein binding, immune system, and oxidation-reduction, while most genes upregulated in aposymbiotic sponges or those fed with bacteria were involved in metabolic processes (fig. 5).

**Figure 5:**
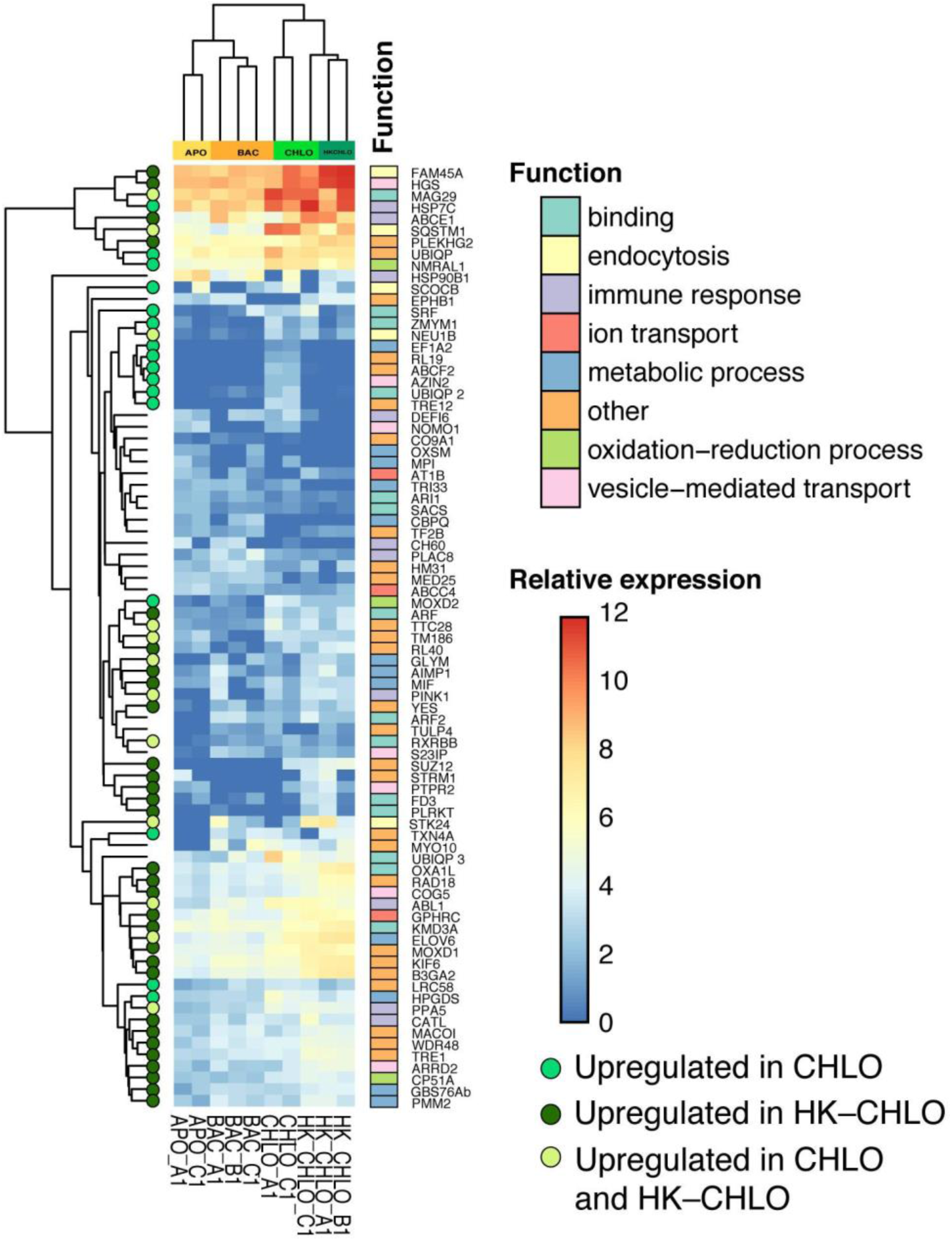
Heat map of subset of differentially expressed genes with BLAST annotations. Heatmap shows significantly (minimum fold change of 2 and a *p*-value of 0.001) differentially expressed genes across four treatments (relative expression colour coding: blue, low expression; red expression, high expression). Aposymbiotic - APO; Bacterial fed - BAC; Heat-killed algal fed - HK–CHLO, and live algal infected - CHLO. Green dots show genes uniquely upregulated in heat-killed algal fed sponges (dark green), live algal infected sponges (medium green) or both (light green). Category functions are provided for genes according to colors on the key to the right.

We compared significant expression differences to identify genes that were regulated in the “digesting” treatments (BAC or HK–CHLO; bacteria or heat-killed algae) compared to live algal infected sponges (CHLO). We asked if there were any genes that were differentially regulated in sponges infected with live native algal symbionts compared to aposymbiotic or bacteria fed sponges and not differentially expressed in heat-killed sponges and found 68 candidate genes to be uniquely upregulated during infection with live algal symbionts (fig. 6A). We considered the genes exclusively up or down regulated in algal infected sponges to be putative “symbiosis related genes” involved in stable residency (Fig 5, fig. 6A).

**Figure 6:**
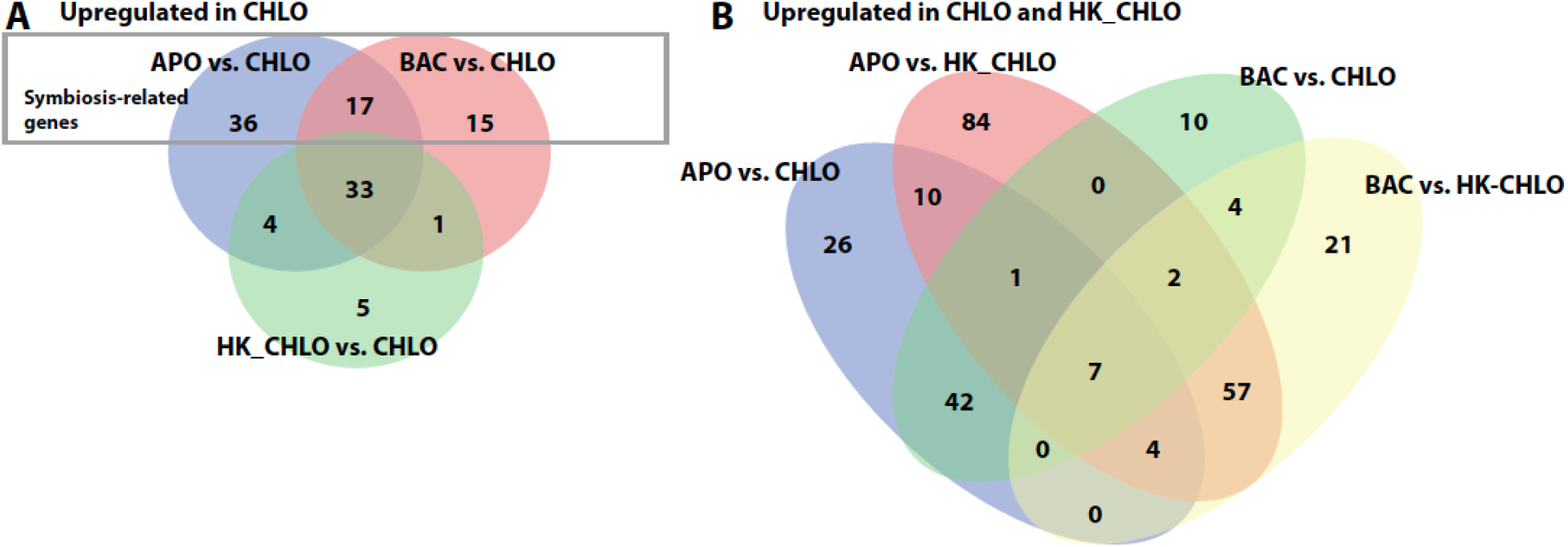
Venn Diagram comparing differentially expressed genes. A. Genes upregulated in live algal infected (CHLO) sponges when compared with Aposymbiotic (APO) sponges, bacterial fed (BAC) sponges, and heat-killed algal fed (HK–CHLO) sponges. B. Genes upregulated in either live algal infected (CHLO), heat-killed fed (HK– CHLO) sponges or both.

Among this set of differentially regulated genes (p<0.001) that are unique to algal infected sponges (CHLO), are some transcripts that we previously found to be significantly differentially regulated at 24 hours post infection in symbiotic compared to aposymbiotic *E. muelleri* (Hall et al. 2021). Genes significantly upregulated at both 4hr and 24hr post infection include a *DBH-like monooxygenase (MOXD2), thioredoxin 4 (TXN4A), leucine-rich repeat containing 58 (LRC58)*, a *glutathione S-transferase (Hematopoietic prostaglandin D synthase, HPGDS)*, and an *NmrA-like family domain-containing protein 1 (NMRL1)* (figs. 5, 7, supplementary table 2). The *MOXD2, TXN4A,* and *NMRL1* may be involved in cellular oxidative-reductive systems. For example, in humans, the *NmrA-like family domain-containing protein 1* (*NMRL1*) functions as a redox sensor that is responsive to NADPH/NADP^+^ levels and helps moderate the production of nitric oxide and reduce apoptosis (Zhao et al. 2008). Indeed, when the photosynthesis of the algae starts, Reactive Oxygen Species (ROS) are released, with the potential of damaging host cell elements (Hamada et al. 2018). Therefore, an appropriate oxidative stress response is usually deployed by the host, as it occurs in the symbiotic partnership between *Chlorella* and *Hydra* (Hamada et al. 2018). But *NMRL1* is also known to negatively regulate the activity of *NF-kappaB* (Gan et al. 2009), a protein whose negative regulation has been implicated in the host response to symbionts in corals (Weis, 2019). The finding that multiple DBH-like monooxygenases were also regulated at 24 hours post infection (Hall et al. 2021) points to a possible role for these enzymes in the onset of algal symbiosis or to a more generalized role of reactive oxygen species signaling in phagocytosis events. Indeed, the finding that multiple glutathione S-transferase genes are upregulated at 24 hours post infection (Hall et al. 2021), along with the up-regulated *glutathione S-transferase* found here, suggests that this gene family may be important for cellular detoxification or for promoting cell growth and viability during symbiosis.

**Figure 7:**
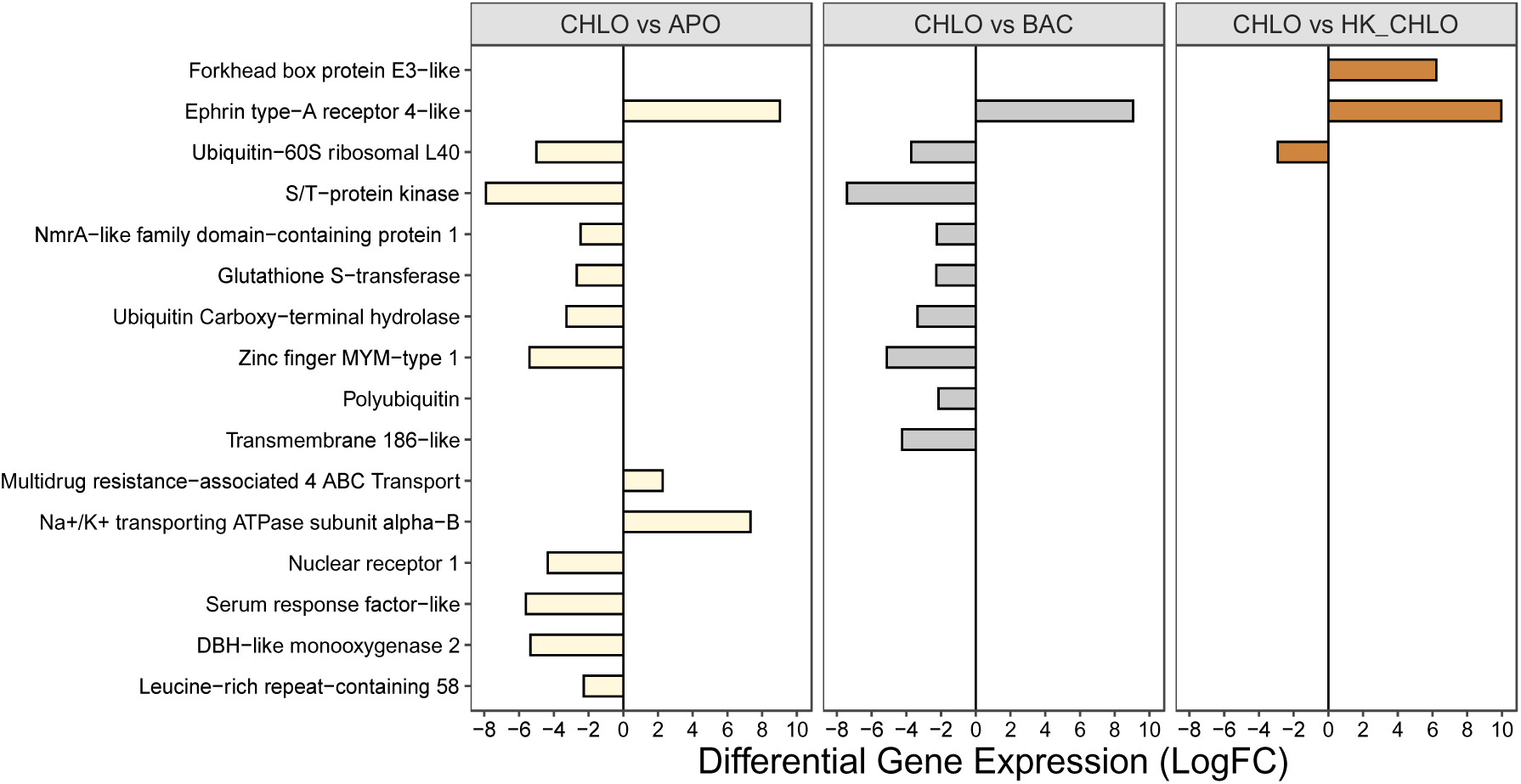
Symbiosis-specific differentially expressed genes. Log fold changes in expression (p<0.001) when comparing live algal infected sponges (CHLO) to each of the other three treatments of algal-free “aposymbiotic” (APO), bacterial-fed (BAC), and heat-killed fed (HK–CHLO). Genes with increased expression in live algal infected sponges compared to given treatments have negative LogFC values and are shown to the left of the solid line on the graph. Only genes with clear BLAST annotations are shown.

Several genes that were uniquely up-regulated at significant levels in the 4 hour post infection treatment were also found in our 24 hour algal symbiosis reference transcriptome, but not regulated at statistically significant levels at the 24 hours post infection timepoint. One of these genes is a *MADS box-containing serum response factor* (*Serum response factor-like*, figs. 5, 7 and supplementary Table 2). These genes have been implicated as Immediate Early Genes (IEGs) that can be activated in response to intrinsic and extrinsic cellular signals and be transcribed rapidly after stimulation. This occurs without the need for *de novo* protein synthesis and often in response to cellular stress (Bahrami & Drablos 2016). Also included in the upregulated genes is a *Nuclear receptor* (*NR1*) or *Retinoic acid receptor RXR-beta-B* (*RXRBB*) gene (figs. 5, 7 and supplementary table 2) containing both a DNA-binding domain as well as a ligand-binding domain of nuclear receptors. These genes are known to function as transcriptional regulators involved in homeostasis, metabolism, and development (Hwang et al. 2014) and can directly translate the message of signaling molecules into a transcriptional response (Miglioli et al. 2021). It seems possible that these host genes may play a role at the onset of symbiosis, but further studies will be needed to evaluate their functions in this biological context.

The upregulation of a probable *ubiquitin carboxy-terminal hydrolase* (*FAM188b*) in algal fed sponges as compared to aposymbiotic and bacterial fed sponges is notable (figs. 5, 7 and supplementary table 2). This enzyme likely has both ligase and hydrolase activity to recycle and remove ubiquitin from degraded proteins as well as linking molecules of ubiquitin for tagging proteins to be discarded. There are many documented roles for these types of proteins, mostly having to do with assessment of cellular ubiquitin and glutathione levels, and regulation of cell cycles. Most interestingly, in regards to the expression trends we see here, is the implication that ubiquitin carboxy-terminal hydrolase can suppress activation of NF-kappaB and increase cellular reactive oxygen species, indicating a possible role in oxidative stress and immune reaction (reviewed in Matuszczak et al., 2020). This protein is also known to play roles in regulation of apoptosis and extrinsic apoptotic signaling and immune defense response. Coupled to this finding, we see upregulation of two ubiquitin genes (likely *polyubiquitin* and *ubiquitin-60S ribosomal L40 (UBIQP),* figs. 5, 7 and supplementary table 2), both of which are implicated in the peroxisome proliferator-activated receptor (PPAR) signaling pathway which has roles in metabolism, cell proliferation/differentiation, as well as immune cell function (Le Menn & Neels, 2018). This is of particular interest since we find upregulation of a *NR1* gene. The upregulated *NR1* gene is part of a superfamily that includes PPARs, suggesting that these genes may be part of the same pathway activated by infection with algal symbionts.

We find relatively few genes that are downregulated in response to symbiosis at 4 hours post infection (fig. 7). Interestingly, two of the downregulated genes are active membrane transporters. One, a *Na^+^/K^+^ transporting ATPase subunit alpha-1* gene, codes for a P-type cation transport ATPase integral membrane protein that can serve to regulate electrochemical gradients by pumping cations across the membrane using primary active transport (Zhiqin & Langhans, 2015). The other is a primary active transporter of the *multidrug resistance-associated 4 ABC transporter group* (*MCR4* or *ABCC4-like*) which is explored in greater detail as described below. We wonder if the downregulation of these transporters may be involved in a mechanism for initiating phagosome arrest, leading to sustained residency in the host cell. Another gene that is decreased in expression in live algal infected sponges is an *ephrin type-A receptor 4-like* gene (*EPB1A*). These proteins are members of a large family of receptor tyrosine kinases that mediate cellular processes, including components of immune function as well as phagocytosis, by interacting with membrane-bound ephrin ligands (Darling & Lamb, 2019). The expression of *EPB1A* expression was decreased by 9-10 fold in symbiotic sponges as compared to every other treatment (fig. 7). The role of these receptors in immune cell activation, as well as mediating immune cell trafficking and implication as receptors for some viral and other pathogens (e.g., malaria parasite) (Darling & Lamb, 2019), make it an interesting candidate to explore in regard to establishing this mutualism. Furthermore, downregulation of this *ephrin-type-A receptor 4 like* gene in live algal infected sponges is contrasted to the upregulation of a distinct, other ephrin-type receptor-like gene in sponges fed heat-killed algae (discussed below), indicating possible roles for these types of receptors in distinguishing symbionts from food.

### Heat-killed related genes and dysregulation of symbiosis

Phagocytosis is an essential cellular process that typically involves the internalization of particles > 0.5 µm. Phagocytotically-driven processes shape many dynamic cellular operations, including development and the destruction of invading pathogens (Kerr et al. 1972; Ren and Savill 1998; Vaux and Korsmeyer 1999). In the case of the latter, phagocytosis plays a role in linking innate and acquired immunity (Savina and Amigorena 2007). For example, Boulais et al. (2010) compared the proteomes of 39 taxa (from protists to mammals) and identified an ancient core of phagosomal proteins primarily involved in phagotrophy and innate immunity. For some heterotrophic organisms, like bactivorous freshwater sponges, phagocytosis is the primary strategy available to capture food. However, phagocytosis is also the major route of entry for many phototrophic symbionts; phototrophic symbionts may co-opt cellular machinery using strategies that are similar to those used by parasites to invade eukaryotic cells (Schwarz 2008).

Somewhat surprisingly, we found a larger group of genes to be differentially regulated when aposymbiotic sponges were fed heat-killed native algae (inactivated symbiont and potential prey) as compared to those fed live native algal symbionts (figs. 4–6). After removing differentially expressed genes that were similarly regulated in live algal infected sponges (fig. 6B), we find genes uniquely regulated in response to the heat-killed algae (fig. 8). We consider these genes to be candidates for phagocytosis and immunity related pathways that may have subtle but important roles in the creation of stable symbiotic states.

**Figure 8:**
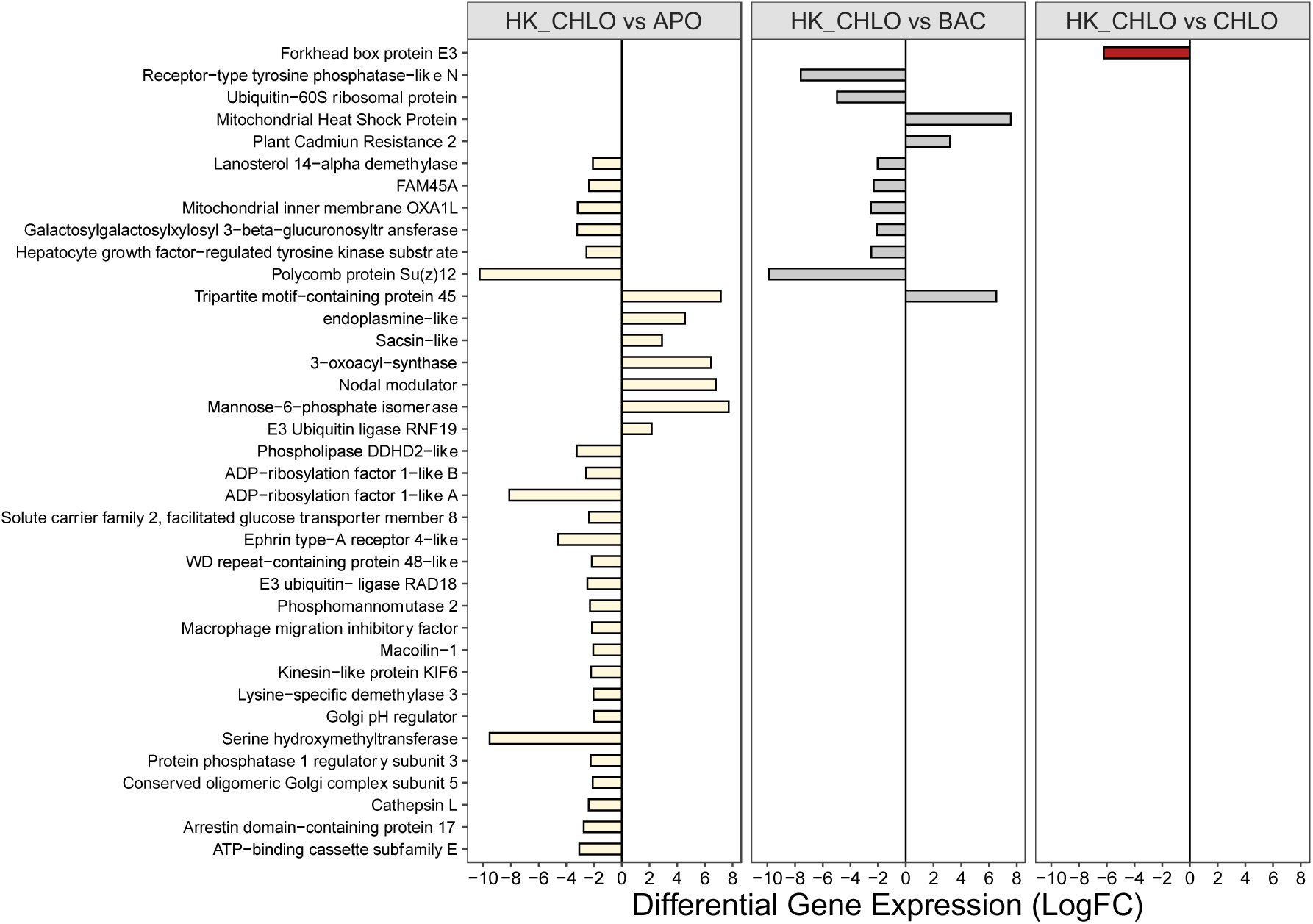
Heat-killed specific differentially expressed genes. Log fold changes in expression (p<0.001) when comparing sponges fed heat-killed algae to each of the other three treatments of algal-free “aposymbiotic” (APO), bacterial-fed (BAC), and live algal infected (CHLO) sponges. Annotated genes with increased expression in heat-killed algae fed sponges compared to given treatments have negative LogFC values and are shown to the left of the solid line on the graph.

Differentially regulated genes with receptor-related activities were identified. One gene that was upregulated when sponges were exposed to heat-killed algae was an *arrestin domain-containing protein*. Arrestins are known to selectively bind to G protein coupled receptors (GPCRs) and other classes of cell surface receptors to control information flow by binding to phosphorylated receptors and blocking further signaling (Chen, Iverson & Gurevich, 2018). For example, the complexes formed between β-arrestin and GPCRs occupied by ligands become targeted for endocytosis and, ultimately, internalization of the receptors (Ferguson et al., 1996). Interestingly, we also find a *hepatocyte growth factor-regulated tyrosine kinase substrate* (*Hrs*) to be increased in expression in sponges fed heat-killed algae (HK–CHLO). These proteins are known to regulate trafficking, degradation, and recycling of GPCRs as part of protein complexes localized on endosomal membranes (Roux et al., 2017). In vertebrate cell lines, *Hrs* associate with phagosomes prior to endosome/lysosome fusion and depletion of *Hrs* impairs the maturation of phagosomes. We find parallels in the infection of cells by mycobacteria - particularly avirulent mycobacteria - which leads to impaired recruitment of *Hrs* to phagosomes and contributes to the arrest of phagosomal maturation (Vieira et al., 2004). The increase in *Hrs* expression here may point to a role for this protein in phagosomal maturation leading to digestion of the heat-killed prey. While we find this gene to be relatively decreased in expression in sponges infected with live algae, that decrease is not significant. Nonetheless, this gene may be a candidate to pursue to determine if there is a role in the arrested phagosome leading to stable algal symbiosis.

We find a *receptor-type tyrosine phosphatase-like N* gene and an *Ephrin type receptor* gene (*tyrosine-protein kinase Yes, YES*) were increased in expression in heat-killed algal treatments (figs. 5 and 8). As discussed earlier, another transcript encoding an *Ephrin type-A receptor 4-like* gene was downregulated in sponges infected with live algae. These proteins are members of a large family of receptor tyrosine kinases that mediate cellular processes, including components of immune function as well as phagocytosis, by interacting with membrane bound ephrin ligands (Darling & Lamb, 2019). Hamada et al (2018) found that an *ephrin type-A receptor* was increased in expression in symbiotic *Hydra* compared to aposymbiotic *Hydra*, but not in a light dependent manner. Activation of these receptors, especially the *Ephrin receptor like* gene, may indicate a role for immune or phagocytic pathways in response to the heat-killed algae that are suppressed during symbiosis and activated during digestion of prey.

A variety of genes that play roles in endocytosis and lysosomal processes are differentially expressed in the heat-killed algal fed treatment (HK-CHLO; figs. 5 and 8). A *conserved oligomeric Golgi complex subunit 5* gene (*COG5*), involved in intra-Golgi vesicle mediated transport, as well as a *Golgi pH regulator* gene (*GPHRC*) are upregulated in the sponge in response to being fed heat-killed algae. Golgi pH regulators are anion channels that are required for acidification and other Golgi functions (Maeda et al., 2008). Also upregulated are two transcripts encoding *ADP-ribosylation factor 1-like* genes (*ARF* and *ARF2*), which are members of the Ras superfamily and play roles in post-Golgi membrane trafficking (Shin et al., 2005), and a *FAM45A* gene, which is involved in regulation of early to late endosomal transport and is a guanine nucleotide exchange factor for Rab small GTPases (Zhang et al., 2019). The upregulation of these genes lend support to the hypothesis that digestive processes, such as the engulfment of the heat-killed algae via endocytosis and digestion via lysosomal fusion, are uniquely upregulated in the heat-killed algae treatment and suppressed in the live-algae treatment.

A large number of genes that are differentially expressed in the heat-killed only treatment are implicated in metabolic processes. This includes *Protein phosphatase 1 regulatory subunit 3* (or *Glycogen-binding subunit 76A*, *GBS76A*), *serine hydroxymethylase* (*GLYM*), *3-oxoacyl-synthase* (*OXSM*), *Phosphomannomutase 2* (*PMM2*), *Mannose-6-phosphate isomerase* (*MPI*), as well as genes like *macrophage migration inhibitory factor* (*MIF*) that play roles in tyrosine and phenylalanine metabolism. Nodal modulator, which is involved in inositol metabolism, is also differentially expressed (figs. 5 and 8). While *MIF* also has many roles in mediating innate immune response (Calandra & Roger, 2003) and nodal modulator, a conserved type1 transmembrane glycoprotein, is involved in structural morphology of the endoplasmic reticulum (Amaya et al., 2021), it is unclear why they are differentially regulated in this particular treatment if not because of their roles in metabolism.

Finally, among the sponge genes upregulated in response to heat-killed algae (figs. 5 and 8), *ATP-binding cassette subfamily E* (*ABCE1*) is known to act as an inhibitor to block the activity of ribonuclease L. Ribonuclease L inhibits protein synthesis involving interferons, which are released during a viral infection to alert nearby cells to increase immune activity. As such, expression of *ABCE1* can promote activity leading to an increase in antiviral defense mechanisms. While there is much work to be done to examine the roles of differentially regulated sponge genes in response to heat-killed algae, it seems clear that many of these genes are candidates for phagocytosis pathways related to the digestion of food particles in freshwater sponges.

### Genes regulated in live and heat-killed algal treatments are enriched for immune function roles

We hypothesized that comparing sponges fed heat-killed algae with those fed live algae might allow us to identify genes with altered expression in both treatments that result as a response to algal specific epitopes (figs. 5, 6, and 9). These genes may be candidates for understanding species-specific recognition pathways or common pathways in initiation of phagocytosis, but not those responsible for regulating stable host:symbiont interaction and persistence of symbionts in the intracellular environment.

The protein *cathepsin L*, widely known for roles in innate immunity as an endolysosomal protease, is upregulated in response to both heat-killed and live algae. At 4 hr post-infection with live algae, *cathepsin L* expression is increased, but not at a statistically significant level; however, by 24 hr post-infection, this transcript is significantly upregulated in response to live algal symbionts (Hall et al., 2021). It is also significantly upregulated at 4 hr post feeding with heat-killed algae (fig. 5, supplementary table 2). This is interesting because *cathepsin L* is also upregulated in response to *Vibrio* spp. in several mollusks and in the freshwater prawn in response to bacterial and viral infections. It plays a role in cell death in the superficial light-organ tissue in *Euprymna scolopes* colonized by *Vibrio fischeri* at the onset of symbiosis (reviewed in Peyer, Kremer & McFall-Ngai 2018). *Cathepsin L* plays numerous roles in immune function, including phagocytosis and non-oxidative killing of bacteria (Muller et al., 2013), and at least one obligate intracellular pathogen is known to manipulate *cathepsin L* activity as a strategy to alter neutrophil function so that the pathogen maintains infection and can proliferate inside the host cell (e.g., Thomas, Samanta & Fikrig, 2008).

The other genes that were regulated in both live algal and heat-killed algal fed sponges are enriched for roles in immune function in other organisms (figs. 5 and 9). *Tyrosine protein kinase Lyn* (*ABL1*), upregulated in live and heat-killed treatments (fig. 5), is often involved in regulation of signaling pathways by phosphorylating inhibitory receptors resulting in modulation of immune cell activation (Xu et al., 2005). *Tartrate-resistant acid phosphatase 5* (*PPA5*) is a lysosomal metalloenzyme expressed in macrophages that promotes inflammation and immune cell activity and is involved in gene ontologies such as negative regulation of interleukin production, negative regulation of tumor necrosis factor production and negative regulation of nitric oxide biosynthetic process (The UniProt Consortium, 2021). Both of these genes continue to be upregulated at 24hr post infection in this symbiosis (Hall et al., 2021).

**Figure 9:**
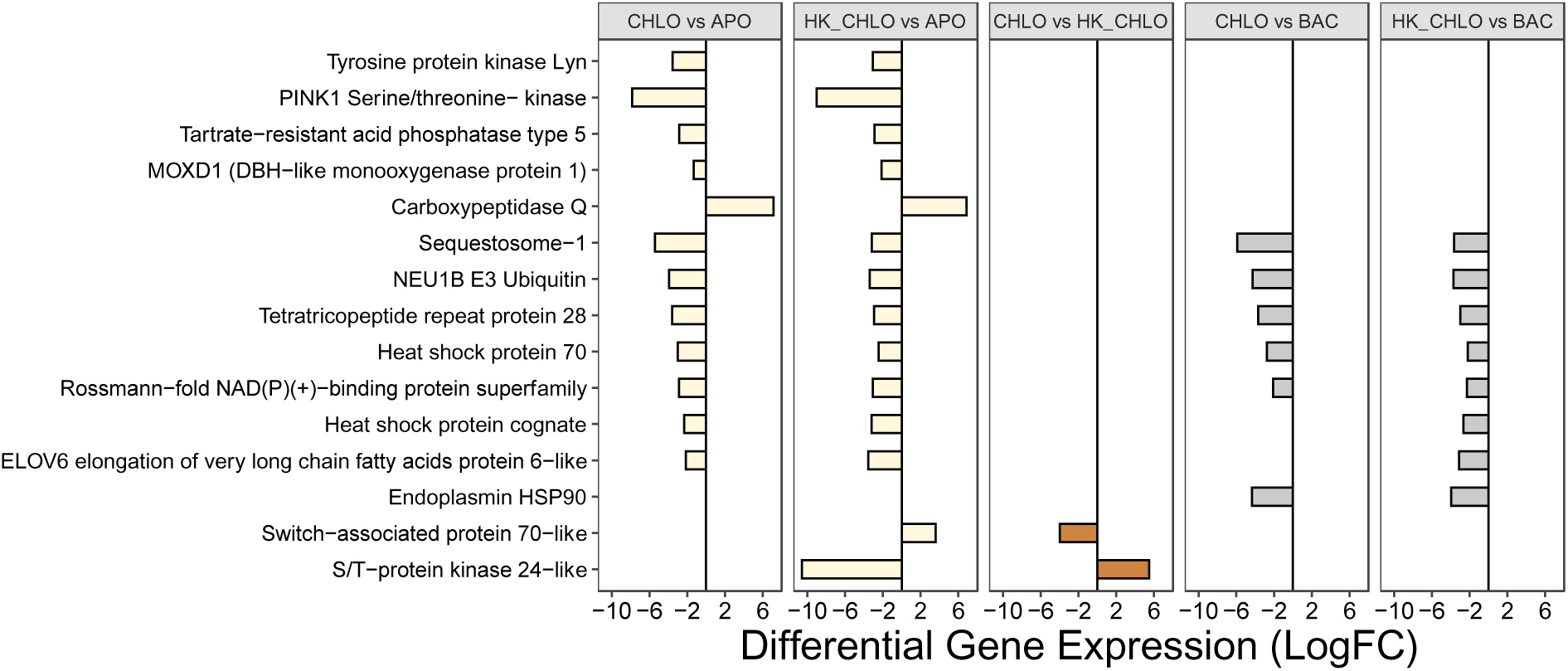
Algal specific differentially expressed genes. Log fold changes in expression (p<0.001) when comparing sponges fed heat-killed algae and live algae to each of the other three treatments of algal-free “aposymbiotic” (APO), bacterial-fed (BAC), and live algal infected (CHLO) sponges. Annotated genes with increased expression in heat-killed algae or live algal fed sponges compared to given treatments have negative LogFC values and are shown to the left of the solid line on the graph.

*PINK1*, a *serine/threonine kinase*, is upregulated in heat-killed and algal fed sponges (figs. 5 and 9). This gene plays roles in protecting cells from stress-induced mitochondrial dysfunction by setting the stage for mitophagy as part of the innate immune response (Song, Zhou & Zhou, 2020). *PINK1* has a wide variety of functions in human systems including regulation of cell metabolism, cancer development and inflammation as well as being responsible for the pathogenesis of early-onset Parkinson’s disease. Recently, *PINK1* was shown to modulate antiviral immune response through the regulation of *TNF receptor-associated factor 3* (Zhou, 2019), another gene we find to be upregulated in algal infected sponges (Hall et al, 2021). We also find *Sequestosome-1* (*SQSTM1*), an autophagy receptor also linked to Parkinson’s disease, to be upregulated in heat-killed and algal infected sponges (figs. 5 and 9). This gene may be involved in regulating the activation of NF-kappaB as well as cell differentiation (The UniProt Consortium, 2021) and is involved in isolation of cargos degraded by autophagy, induction of antioxidant responses, and regulation of inflammation, apoptosis and endosomal trafficking. Interestingly, *sequestosome-1* also plays important roles during *PINK1* mediated mitophagy and is linked with some E3 ubiquitin ligases for its roles in ubiquitylation, autophagy, and recruitment of endocytotic vesicles (reviewed in Sánchez-Martín & Komatsu, 2018). Of note, we do find a *E3 Ubiquitin Protein Ligase 1B-like* gene (*NEU1B*) among the upregulated transcripts in our heat-killed and live algal fed sponge treatments (figs. 5 and 9). This may indicate conserved connections between these immune pathway regulators that have yet to be elucidated.

The upregulation of expression of several heat shock proteins in live and heat killed algal-fed sponges is not surprising (figs. 5 and 9), as heat shock proteins play crucial roles as components of cellular response to stress, modulation of phagocytosis and immune function (e.g., reviewed in Lee& Repasky, 2012; Zininga, Ramatsui, & Shonhai, 2018). Here, we find a *heat shock protein 70* (*HSP70*), known to stimulate phagocytosis of internalized antigens in macrophages (Wang et al., 2006), a heat shock protein cognate (*Heat shock cognate 71 kDa*, *HSP7C*), which can mediate LPS-induced inflammatory responses through modulation of NF-kappaB in macrophages (Sulistyowati et al., 2018), and *endoplasmin HSP90* (*HSP90B1*), which is an essential immune chaperone acting to fold proteins in secretory pathways and protect the integrity of associated antigens (Calderwood, Gong, & Murshid, 2016) all to be differentially expressed in response to algae. The role of heat shock proteins have been explored in coral symbioses, mostly in relation to thermal stress and bleaching (e.g., Ishii et al., 2019; Ross 2014; Leggat et al., 2011), but it is known from this literature that both algae and coral employ heat shock proteins during stress response (Weis 2008). The role of these proteins in modulating sponge immune responses or phagocytosis remains to be explored.

Finally, we find a *switch-associated protein 70-like* gene (*Differentially expressed in FDCP 6 homolog*, *DEFI6*) to be upregulated in live and heat-killed algal fed sponges (figs. 5 and 9). This phosphatidylinositol 3,4,5-trisphosphate-dependent guanine nucleotide exchange factor is known to be recruited to phagosomes, to transduce signals from tyrosine kinase receptors to Rac, and to mediate signaling of membrane ruffling by organization of actin cytoskeleton as an essential process for phagocytosis (Baranov et al., 2016). The differential regulation of the genes described above in live and heat-killed algal treatments here show that regardless of whether sponge cells will form stable symbioses with live algae or digest heat-killed algae, a variety of putative immune and phagocytosis response pathways are activated before the phagosome arrests or digests algae.

### ABC Transporter suppression during onset of symbiosis

ABC (ATP-Binding Cassette) transporters encompass the largest family of transmembrane proteins and originated in the first unicellular organisms (Rees, Johnson & Lewinson, 2009). Using energy released by ATP hydrolysis, these transport proteins are responsible for the movement of ions and other molecules into the cell (influx) and out of the cell (efflux), and are respectively classified as importers and exporters (Vasiliou, Vasiliou & Nebert, 2009). The ABCC subfamily consists of transporters that have a diverse functional spectrum, including ion transport, acting as cell surface receptors, and secretion of toxins (Dean, Hamon & Chimini, 2001). One distinct feature that many ABCC transporters possess that distinguish them from other ABC transporters is they are responsible for the transportation of glutathione conjugates or cotransport glutathione combined with a substrate (Stolarczyk, Reiling & Paumi, 2011).

While there is limited evidence describing the role of ABC transporters in animal symbiotic relationships, many plant-bacteria interactions are facilitated by the host’s ABC transporters (Zhang, Blaylock & Harrison, 2010) and, in animals, there is evidence of downregulated ABC transporter expression as a result of numerous types of infections (Hinoshita et al., 2001). In rats, an increase in a cytokine immune response results in a decrease of expression of ABC transporters and one ABC transporter (*ABCC2*) is downregulated by NF-κB (Nakamura et al., 1999). We previously showed that a *TNF receptor-associated factor* in the NF-κB signaling pathway was found to be significantly increased in expression at 24 hours post infection with live native algae (Hall et al., 2021). This *TNF receptor associated factor* is also known to be directly involved in cytokine production, which could potentially downregulate certain ABC transporters in *E. muelleri* as is suggested to occur in other model systems.

The *E. muelleri ABCC4* gene (scaffold Em0024g219) is 10,820 base pairs long and consists of 29 exons and 30 introns. There are multiple domains that identify it as a predicted ABCC transporter, including the Multi drug resistance-associated protein (MRP) domain with its highly conserved C-terminal region, the PLN3130 and ATP-binding cassette domain found on the nucleotide-binding (NBD) domains, and the Md1B domain encoding for the ATPase and permease component of the transporter. Included is the transmembrane (TMD) region of the protein that is composed of six transmembrane helices, thus indicating that a TMD is present. The AAA domain also codes for an ATPase. Both of the ATPases encoded by separate domains are located in the NBD subunit, which is responsible for the hydrolysis of ATP to ADP. The N-terminal helix, which is poorly conserved across species, is not present in this sequence.

*E. muelleri ABCC4* expression was downregulated in the sponges fed live algae (fig. 5) 4hr after infection, but was further evaluated using qRT-PCR at four post-infection time points and compared to expression levels of aposymbiotic sponges cultured for the same time period under identical experimental conditions (fig. 10). It appears that after the initial downregulation of *ABCC4* expression following symbiont introduction at 4 hours post infection, the levels of expression continued to decrease at 8 hours post infection and reached a minimum at 24 hours post infection. When measured at 48 hours post infection, the levels of *ABCC4* expression began to increase again back to aposymbiotic levels. This trend was consistent when the data was normalized with *GAPDH* and *Ef1a* housekeeping genes.

**Figure 10:**
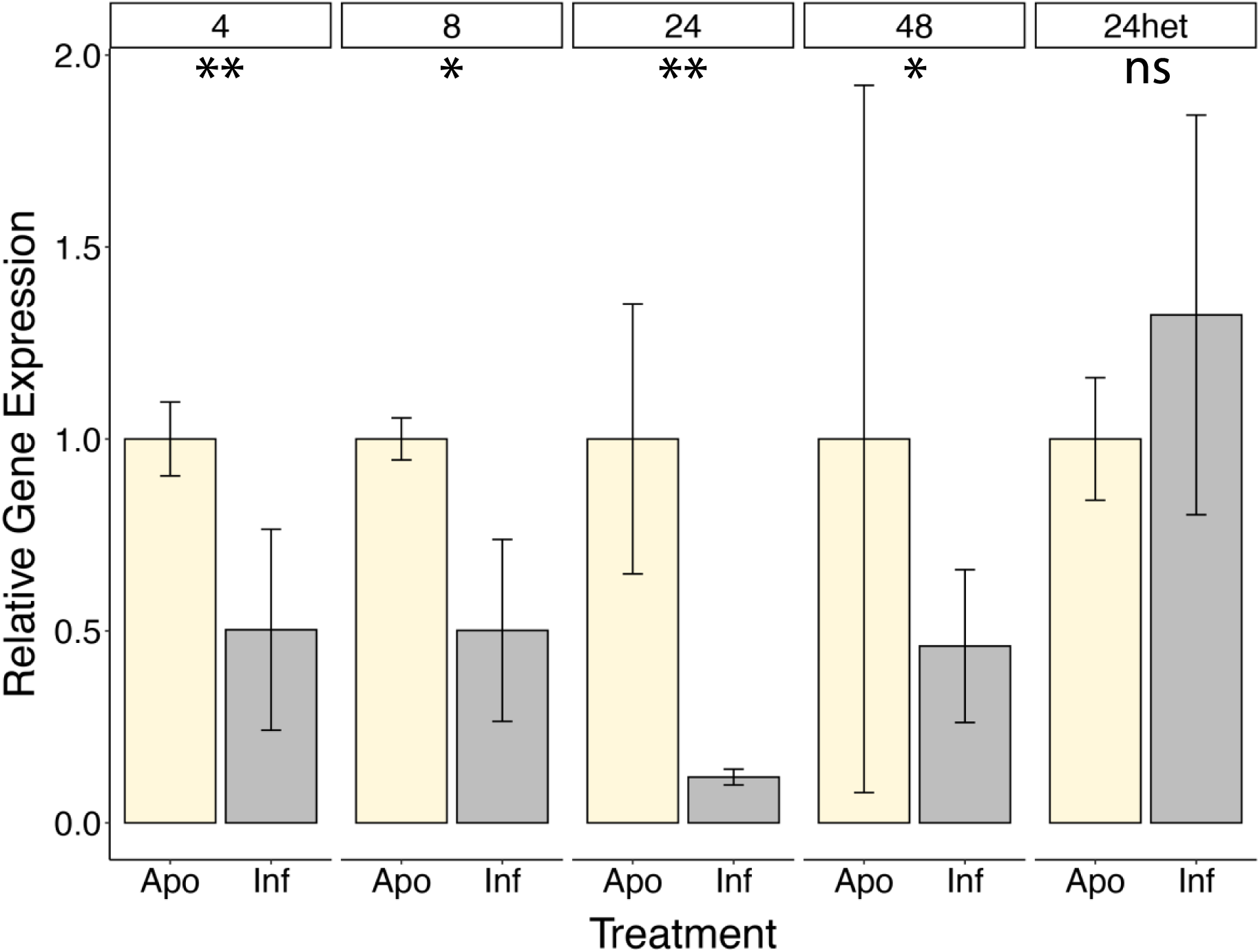
Differential *ABCC4* expression. Relative gene expression for *ABCC4* is shown for five different experiments for sponges infected with live algal symbionts relative to aposymbiotic age-matched sponges. Sponges infected with native symbionts at 4, 8, 24, and 48 hours post infection or with heterologous symbionts at 24 hours post infection are given. Error bars represent SD. 2-tailed, two sample independent t-tests were performed on deltaCt values (** = p<0.001; * = p<0.05; ns = not significant).

To determine if the downregulation of *ABCC4* in *E. muelleri* was a generalized response to live algal symbionts, or specific to native algal infection, we infected sponges hatched from gemmules collected from Bryant Park, VA with native algae and with *Chlorella*-like algae isolated from *E. muelleri* sponges collected from Lake O’Connor, Canada (provided by Dr. Sally Leys). Relative levels of gene expression for *E. muelleri ABCC4* was evaluated for the heterologous cross infections. For Bryant Park sponges infected with algae isolated from Lake O’Connor *E. muelleri*, *ABCC4* expression levels increased, but not significantly, when compared to aposymbiotic Bryant Park sponges. The change in *ABCC4* expression in the infected sponges was not statistically significant with either housekeeping gene used for normalization. This data suggests that the geographic variant did not elicit the same response (downregulation of *ABCC4*) as the native algae and that these sponges may form symbiotic relationships with closely related, but distinct algal populations.

## CONCLUSIONS

Much work will be needed to understand the nature of these facultative phototroph:heterotroph partnerships in terms of how stable residencies within host cells is achieved, but the possibilities provided by *E. muelleri* and its algal symbionts as a model for studying symbiotic relationships is promising. We have identified a suite of genes that are regulated early on during the establishment of a stable symbiosis between *E. muelleri* and its native green algal symbionts. We have also begun to differentiate these genes from those involved in generalized phagocytosis events related to feeding and/or immunity. While the majority of differentially regulated genes are proteins of uncharacterized or unknown function, the genes of known function point to pathways that play interesting roles in establishing symbiotic states. These include cellular oxidative-reductive systems, response to cellular signals, membrane and vesicle–mediated transport, metabolism, cell proliferation/differentiation, as well as immune cell function.

## METHODS

### Sample collection

Sponge gemmules were collected and processed as described in Hall et al. (2021). Sponges for this study were either collected in Richmond, VA in Bryan Park (37°35’53.0’’ N, 77°28’06.3’’W) under Virginia Department of Game and Inland Fisheries Permit #047944 or were obtained from Dr. Sally Leys (Alberta) from a site at O’Connor Lake, Saskatchewan, Canada (61.3129° N, 111.8608° W). Gemmules were isolated, washed and stored as described in Leys, Grombacher & Hill (2019).

Native sponge algae were isolated as described in Hill, Nguyen & Hill (2020). In the case of Bryan Park *E. muelleri*, the algae were isolated from adult tissue (Hall et al. 2021) and from *E. muelleri* hatched gemmules in the case of Lake O’Connor sponge-derived algae. Algal cultures were propagated and used in subsequent reinfection experiments and grown at ± 22°C under fluorescent light for 16 hour per day in either Basal Medium (Sigma-Aldrich, Milwaukee, WI) or in Modified Bolds 3N Medium (UTEX, Austin, TX).

### Algal infection of sponges

Gemmules of *E. muelleri* were used from stocks that were either frozen at −80°C or held at 4°C. Gemmules were hatched and cultured as described in Leys, Grombacher and Hill (2019) and infections were initiated once sponges had a functioning canal system and were pumping water through their tissues. Live sponge-derived algal cells (“CHLO” treatment), algal cells that had been heat-killed by boiling for 3 min (“HK–CHLO” treatment), or living non-pathogenic K12 *E. coli* bacteria (“BAC” treatment) were introduced into the water surrounding the sponge. Infections were conducted as described in Hill, Nguyen and Hill (2020). Infections were initiated with 130,000 cells ml^-1^1X Strekal’s harvested during the logarithmic portion of their growth phase. We estimated cell densities and population growth characteristics using optical density (OD) measurements at 425 nm and 675 nm for algae and 600 nm for bacteria. Algae or bacteria were slowly pipetted around and above the tissue to inoculate sponges and placed under fluorescent grow lights at room temperature for 4 hours. Infected sponges used at later time points were placed under a 12:12 light:dark exposure.

### Microscopy

Sponges were grown on 35 mm glass bottom dishes (MatTek Life Sciences) and either infected with live sponge-derived algae or left untreated. Fixation and imaging were as described in Hall et al. (2021). Briefly, sponges were fixed in 4% paraformaldehyde and 1/4 Holtfreter’s Solution overnight at 4°C. After washing and permeabilization, tissue was stained with Hoescht 33342 (1:200 dilution, Thermo Fisher Scientific, Waltham, MA) and Phalloidin Alexa 488 (1:40 dilution, Thermo Fisher Scientific, Waltham, MA) and imaged using an Olympus FV1200 laser scanning microscope using FluoView software.

Electron microscopy was performed as described in Hall et al. (2021). Briefly, sponge samples infected with live algae were fixed in 2.5% glutaraldehyde, washed in 0.2 M cacodylate buffer (pH 7.4) and postfixed with 1% OsO4 and 1% uranyl acetate.Samples were dehydrated, infiltrated in propylene oxide, and embedded in Embed 812 plastic resin. Ultrathin sections were stained with uranyl acetate and lead citrate. Micrographs were taken using a JEOL 1010 transmission electron microscope at the University of Richmond with an Advanced Microscopy Techniques XR-100 Digital CCD system.

### RNA isolation for library construction, and sequencing

Sponges were grown from gemmules in 1X Strekal’s to the stage where a functioning osculum had developed. To triplicate samples of these sponges (∼20-30 sponges per treatment), we added live algal cells (CHLO), heat-killed algal cells (HK–CHLO), or bacteria (BAC) as described above. In addition, a batch of sponges received no treatment and were considered the control (APO). Tissue was collected after 4 hours of exposure to algae, heat-killed algae, bacteria or no exposure and washed several times to remove algae or bacteria from the surrounding water and surfaces, and stored at - 80°C after RNA*later* treatment (Thermo Fisher Scientific, Waltham, MA). Total RNA was isolated from 3 replicates of each of the four treatments using TRIzol™ Reagent (ThermoFisher Scientific) and the standard protocol according to the guidelines of the manufacturer. Then, mRNA purification was performed with Dynabeads mRNA DIRECT kit (ThermoFisher Scientific), only performing the final stage of the protocol, ‘Elimination of rRNA contamination’ with the purified total RNA as input. The quantity and overall quality of mRNA were assessed on a NanoDrop 2000 (ThermoFisher Scientific), and only those extractions with A260/280 with values around 2 were used. The cDNA synthesis for the libraries was performed with Scriptseq v2 kit (Illumina) (according to the manufacturer’s instructions), using an initial mRNA quantity of 50 ng. For the final cDNA libraries, we selected a fragment size of approximately 250-350 bp with a BluePippin (Sage Science, MA, USA). The final concentration of the cDNA libraries was assessed with a Qubit™ dsDNA HS Assay kit (ThermoFisher Scientific) and the quality with an Agilent Tapestation 2200 system (Agilent Technologies). The sequencing of all libraries was done in several runs in an Illumina NextSeq 500 platform at the Natural History Museum of London sequencing facility (Molecular Core Labs) with 150 bp paired-end settings.

### Transcript assembly and analysis

The raw reads were filtered based on quality with Trimmomatic (Bolger et al., 2014), cutting when the average quality in a 4-base window dropped below 28. Then, high quality reads were used for *de novo* assembly, which was done with Trinity v2.8.4 (Grabherr et al., 2011). Completeness of the assembly was calculated with Benchmarking Universal Single-Copy Orthologs (Busco V2/3) against metazoan and eukaryotic cassettes (Simão et al., 2015) in gVolante (Nishimura et al. 2017).

To obtain the gene expression metrics associated to each treatment, we mapped the reads to the reference assembly with Bowtie2 (Langmead and Salzberg, 2012), performed transcript quantification with RSEM (Li and Dewey, 2011), and then analysed differential gene expression (DGE) analysis with edgeR (Robinson et al., 2009; McCarthy et al., 2012) using a minimum fold change of 2 and a *p*-value of 0.001. We analysed the differences in gene expression in pairwise comparisons between all four treatments.

To annotate the transcriptome, we first did a blastx search (Altschul et al., 1997) of the transcriptome using the *refseq* database (Pruitt et al., 2007) (accessed in 2018) with a cut-off e-value of 1e-5. The sequences with blast hits were visualized in MEGAN (Huson et al. 2007) for taxon assignment and further annotated by Blast2GOPRO (Conesa et al., 2005) to retrieve the functional information from the Gene Ontology (GO) terms.

### RNA Isolation, cDNA synthesis and differential expression analysis by qRT-PCR

Sponge tissue from time points post-algal infection or from control sponges was harvested by scraping tissue from the petri dish using a pipet and placed in RNA*later* solution (Thermo Fisher Scientific, Waltham, MA). Per biological replicate, sponges were pooled as 12-24 samples per time point. RNA was isolated from tissue samples using the Aurum Total RNA Mini Kit (Qiagen, Hilden, Germany) using the Spin Format in biological triplicate according to the manufacturer’s instructions. Additional on-column DNase I steps were performed to limit DNA contamination. RNA quantity was determined using a Nanodrop Spectrophotometer. If sponge tissue was not immediately processed into RNA, it was placed in RNAlater and stored at 4°C. RNAlater was removed after 24 hours and tissue was stored at −80℃ until use.

For each biological triplicate, cDNA was synthesized using the AffinityScript qPCR cDNA Synthesis Kit (Agilent Technologies, Santa Clara, CA) according to the manufacturer’s instructions. Starting RNA sample concentration ranged depending on the set of experiments, but was typically in the range 125-250 ng/µl. qRT-PCR was performed on a Agilent AriaMX Real-Time PCR System (Agilent Technologies, Santa Clara, CA) and master mixes for each qRT-PCR reaction used PowerUp SYBR Green Master Mix (Thermo Fisher Scientific, Waltham, MA). qRT-PCR was run at the following conditions: 5 °C for 10 minutes, 40 cycles at 95℃ for 30 seconds, 55℃ for 1 minute, and 72℃ for 1 minute, and lastly one cycle at 95℃ for 1 minute, 55℃ for 30 seconds and 95℃ for 30 seconds. Triplicate technical reactions were run for each sample in addition to a non-template control also run in triplicate. Primer sequences for *ATP Binding Cassette Subfamily C Member 4* (*ABCC4*) were; forward-ATG CAT GTC TCG TCT CCT CG, reverse-AAT GCC ACA TTG GTG TCT GC. Both *glyceraldehyde 3-phosphate dehydrogenase* (*GAPDH*) and *elongation factor 1-alpha* (*EF1α*) were run with each gene tested as housekeeping genes. The primer sequences for *GAPDH* were; forward-GGT CGA GTC ACA GGT GTT, reverse-CCA AGC AGT TGG TAG TGCA. The primer sequences for *Ef1α* were forward-GCG GAG GTA TCG ACA AGC GT, reverse-AGC GCA ATC GGC CTG GGA CG. Triplicate Ct values were averaged and the expression of *ABCC4* was normalized to the geometric mean of the reference housekeeping genes. Gene expression was analyzed using a modification to the delta delta-Ct method described by Vandesompele et al (2002) and Hellemans et al (2007) to allow for use of multiple housekeeping genes and for efficiency corrected ΔCT values. Resulting relative quantities on the log_2_ scale were used for statistical analysis (2-tailed, 2 sample independent t-test) using R (version 3.6.3).

## SUPPLEMENTARY MATERIALS

### Supplemental Figures

Supp Fig 1: APO_vs_BAC volcano plot

Supp Fig 2: APO_vs_CHLO volcano plot

Supp Fig 3: APO_vs_HK_CHLO volcano plot

Supp Fig 4: BAC _vs_CHLO volcano plot

Supp Fig 5: BAC_vs_HK_CHLO volcano plot

Supp Fig 6: CHLO_vs_HK_CHLO volcano plot

Supp Fig 7: Principal Components Treatments

### Supplemental Tables

Supplementary Table 1: Number of reads and transcriptome assembly and annotation metrics.

Supplementary Table 2: Differentially expressed genes across treatments using edgeR.

## Acknowledgements

This work was supported by the National Science Foundation (Award #1555440) to April L. Hill and Malcolm S. Hill and by an Institutional Development Award (IDeA) from the National Institute of General Medical Sciences of the National Institutes of Health (#P20GM103423).

